# Association Tests Using <u>**Co**</u>py <u>**N**</u>umber Profile <u>**Cur**</u>ves (CONCUR) Enhances Power in Rare Copy Number Variant Analysis

**DOI:** 10.1101/666875

**Authors:** Amanda Brucker, Wenbin Lu, Rachel Marceau West, Qi-You Yu, Chuhsing Kate Hsiao, Tzu-Hung Hsiao, Ching-Heng Lin, Patrik K. E. Magnusson, Patrick F. Sullivan, Jin P. Szatkiewicz, Tzu-Pin Lu, Jung-Ying Tzeng

## Abstract

Copy number variants (CNVs) are the gain or loss of DNA segments in the genome that can vary in dosage and length. CNVs comprise a large proportion of variation in human genomes and impact health conditions. To detect rare CNV association, kernel-based methods have been shown to be a powerful tool because their flexibility in modeling the aggregate CNV effects, their ability to capture effects from different CNV features, and their ability to accommodate effect heterogeneity. To perform a kernel association test, a CNV locus needs to be defined so that locus-specific effects can be retained during aggregation. However, CNV loci are arbitrarily defined and different locus definitions can lead to different performance depending on the underlying effect patterns. In this work, we develop a new kernel-based test called CONCUR (i.e., <u>Co</u>py <u>N</u>umber profile <u>Cur</u>ve-based association test) that is free from a definition of locus and evaluates CNV-phenotype association by comparing individuals’ copy number profiles across the genomic regions. CONCUR is built on the proposed concepts of “copy number profile curves” to describe the CNV profile of an individual, and the “common area under the curve (cAUC) kernel” to model the multi-feature CNV effects. Compared to existing methods, CONCUR captures the effects of CNV dosage and length, accounts for the continuous nature of copy number values, and accommodates between- and within-locus etiological heterogeneities without the need to define artificial CNV loci as required in current kernel methods. In a variety of simulation settings, CONCUR shows comparable and improved power over existing approaches. Real data analyses suggest that CONCUR is well powered to detect CNV effects in gene pathways associated with phenotypes using data from the Swedish Schizophrenia Study and the Taiwan Biobank.

**Author summary:** Copy number variants comprise a large proportion of variation in human genomes. Large rare CNVs, especially those disrupting genes or changing the dosages of genes, can carry relatively strong risks for neurodevelopmental and neuropsychiatric disorders. Kernel-based association methods have been developed for the analysis of rare CNVs and shown to be a valuable tool. Kernel methods model the collective effect of rare CNVs using flexible kernel functions that capture the characteristics of CNVs and measure CNV similarity of individual pairs. Typically kernels are created by summarizing similarity within an artificially defined “CNV locus” and then collapsing across all loci. In this work, we propose a new kernel-based test, CONCUR, that is based on the CNV location information contained in standard processing of the variants and removes the need for any arbitrarily defined CNV loci. CONCUR quantifies similarity between individual pairs as the common area under their copy number profile curves and is designed to detect CNV dosage, length and dosage-length interaction effects. In simulation studies and real data analysis, we demonstrate the ability of CONCUR test to detect CNV effects under diverse CNV architectures with power and robustness over existing methods.

## Introduction

Copy number variants (CNVs) are unbalanced structural variants that are typically 1 kilobase pair (kb) in size or larger and lead to more or fewer copies of a region of DNA with respect to the reference genome. CNVs are typically characterized by two descriptive features. The first feature is CNV dosage, or the total number of copies present, with > 2 copies corresponding to duplications and < 2 copies corresponding to deletions. The second is the CNV length, typically measured in base pairs (bp) or kilobase pairs.

CNVs are important risk factors for many human diseases and traits, including Crohn’s disease, HIV susceptibility, and body mass index [1–3]. Large and rare CNVs are particularly implicated in neuropsychiatric disorders including autism spectrum disorder, schizophrenia, bipolar disorder, and attention deficit disorder [4]. For example, multiple studies have confirmed a greater burden of rare CNVs in schizophrenia cases compared with normal controls, both genome-wide and in specific neurobiological pathways important to schizophrenia (e.g., calcium channel signaling and binding partners of the fragile X mental retardation protein).

Typically, rare CNVs (e.g., < 1% frequency) in the genome are intractable to test individually for disease association and instead are examined with collapsing methods. Collapsing methods summarize variant characteristics across multiple variants in a targeted region, typically a gene set or the whole genome, and perform a test of the collective CNV effects. By accumulating information across multiple rare variants, collapsing methods can have enhanced power to detect the effects of rare CNVs that are hard to detect individually but collectively have a significant impact. Collapsing tests for rare CNVs are primarily built on the foundation of rare single nucleotide polymorphism (SNP) association tests but with additional complexity to accommodate the length and dosage features of CNVs. As with SNPs, the effects of CNVs can vary between loci, but CNV collapsing tests must also account for within-locus heterogeneity due to differential dosage effects or length effects within a CNV region.

Similar to SNP collapsing tests, there are also two families of tests for rare CNV analysis: burden-based methods and kernel-based methods. Burden-based tests, e.g., Raychaudhuri et al. [5], summarize the CNV features of an individual via the total CNV counts or average length and model the CNV effects as fixed effects assuming etiological homogeneity of features across multiple CNVs of a targeted region. Kernel-based tests, e.g., CCRET [6] and CKAT [7], aggregate CNV information via genetic similarity based on certain CNV features and model CNV effects as random effects to account for the between-locus etiological heterogeneity. By design, burden tests are optimal when the association signal is driven by homogeneous effects across CNVs, and kernel-based tests are optimal in the presence of etiological heterogeneity. Burden tests often need to subset CNVs by dosage (e.g., deletions only or duplications only) or size (e.g. > 100kb, > 500kb) to increase homogeneity while kernel-based tests do not have such requirements.

In this work, we focus on kernel-based methods because etiological heterogeneity is becoming a more practically encountered scenario as high-resolution CNV detection technologies permit the detection of CNVs with smaller length. In kernel-based association tests, the association between CNVs and the trait is evaluated by examining the correlation between trait similarity and CNV similarity quantified in a kernel. For kernel construction, we can refer to kernel-based tests for SNPs; since SNPs are evaluated at the same single base-pair position (referred to as a locus) across individuals, it is natural to assess similarity locus-by-locus and aggregate the locus-level similarity over all loci in the target region to obtain an overall SNP similarity. A locus unit for CNVs, however, is not so obvious since CNVs span multiple base pairs and may overlap partially between individuals.

To address this issue, standard CNV kernel-based tests construct CNV regions (CNVR). For example, the CNV Collapsing Random Effects Test (CCRET) [6] creates CNVR by clustering CNV segments of different individuals with some arbitrary amount of overlap (e.g., 1 base pair overlap, 50% reciprocal overlap). With CNVRs, the CNV similarity between an individual pair can be quantified first within each CNVR, and this CNVR-level similarity can be summed over all CNVRs in the target region to characterize overall CNV similarity. However, a drawback of this approach is that CNVRs defined in this fashion are contingent on the unique CNV overlapping patterns among individuals in a study, and the defined CNVRs can vary from one study to another. The arbitrary choice of overlapping threshold also impacts the formation of locus units and consequently how the “between-locus” and “within-locus” heterogeneous effects of CNVs are accounted for.

To avoid the issues introduced by arbitrarily defined CNVRs as in CCRET, the CNV Kernel Association Test (CKAT) [7] adopts a different strategy to quantify CNV similarity between two individuals. Specifically, CKAT allows users to define the CNVR as a biologically relevant region, e.g., a chromosome. CKAT also introduces a new kernel function to measure CNV similarity based on both dosage and length features between two CNV events. This CNV-level similarity is then aggregated to derive a measure of CNVR-level similarity using a shift-by-one scanning algorithm that “aligns” CNVs in two profiles based on their ordinal position. A multiple-testing correction is applied if multiple CNVRs are involved in the targeted region. Although the new strategy bypasses the need of an arbitrarily defined locus unit, the scanning alignment may yield unreliable results if CNVRs are too large and distant CNVs contribute to an inaccurate model of profile similarity. In addition, there are computational considerations with a scanning algorithm. Furthermore, CKAT aligns pairs of CNVs based on their ordinal position rather than considering all possible pairs which may not optimally capture similarity.

To address these challenges in quantifying CNV similarity using kernel-based methods, in this work we propose a new approach called the <u>Co</u>py <u>N</u>umber profile <u>Cur</u>ve-based (CONCUR) association test. Based on the concept of copy number (CN) profile curves (introduced below), the CONCUR association test naturally incorporates both CNV dosage and length features and can capture their main effects as well as dosage-length interactions. Additionally, building the kernel based on CN profile curves permits the quantification of CNV similarity without the need for pre-specified locus units. Moreover, CNV length may be incorporated flexibly in units which are supported in good resolution by the sequencing technology or which are computationally stable. Like CCRET and CKAT, the test is built in the framework of kernel machine regression and is powerful under heterogeneous signals and can adjust for confounders. In this analysis, we use simulation studies to demonstrate the improved power CONCUR over existing kernel-based methods in a variety of settings and illustrate the practical utility of CONCUR by conducting pathway analysis on the Swedish Schizophrenia Study data and the Taiwan Biobank data.

## Results

### Overview of CONCUR

CONCUR assesses the collective effects of rare CNVs on a phenotype in a kernel machine regression framework where the kernel construction does not require a defined CNV locus. As such, CONCUR is built on two major components: (a) the CN profile curve, with which we describe an individual’s CNVs across the genome or a region of interest; and (b) the common area under the curve (cAUC) kernel, with which we measure CNV similarity between two individuals and characterize the CNV effects on the phenotype. In a CN profile curve (e.g., Fig 1), CNV dosage is shown on the y-axis as jumps or troughs diverging from a baseline of 2 copies; the start and end points of the jumps and troughs correspond to the start and end locations of the CNV and are shown on the x-axis. At genomic locations where there are no CNV events, the y-axis (dosage) takes value 2 (i.e., the baseline value). CN profile curves are intended to be a visualization of CNV activity and concurrence across samples and contribute to the CONCUR method through the concept of cAUC.

**Fig 1.**
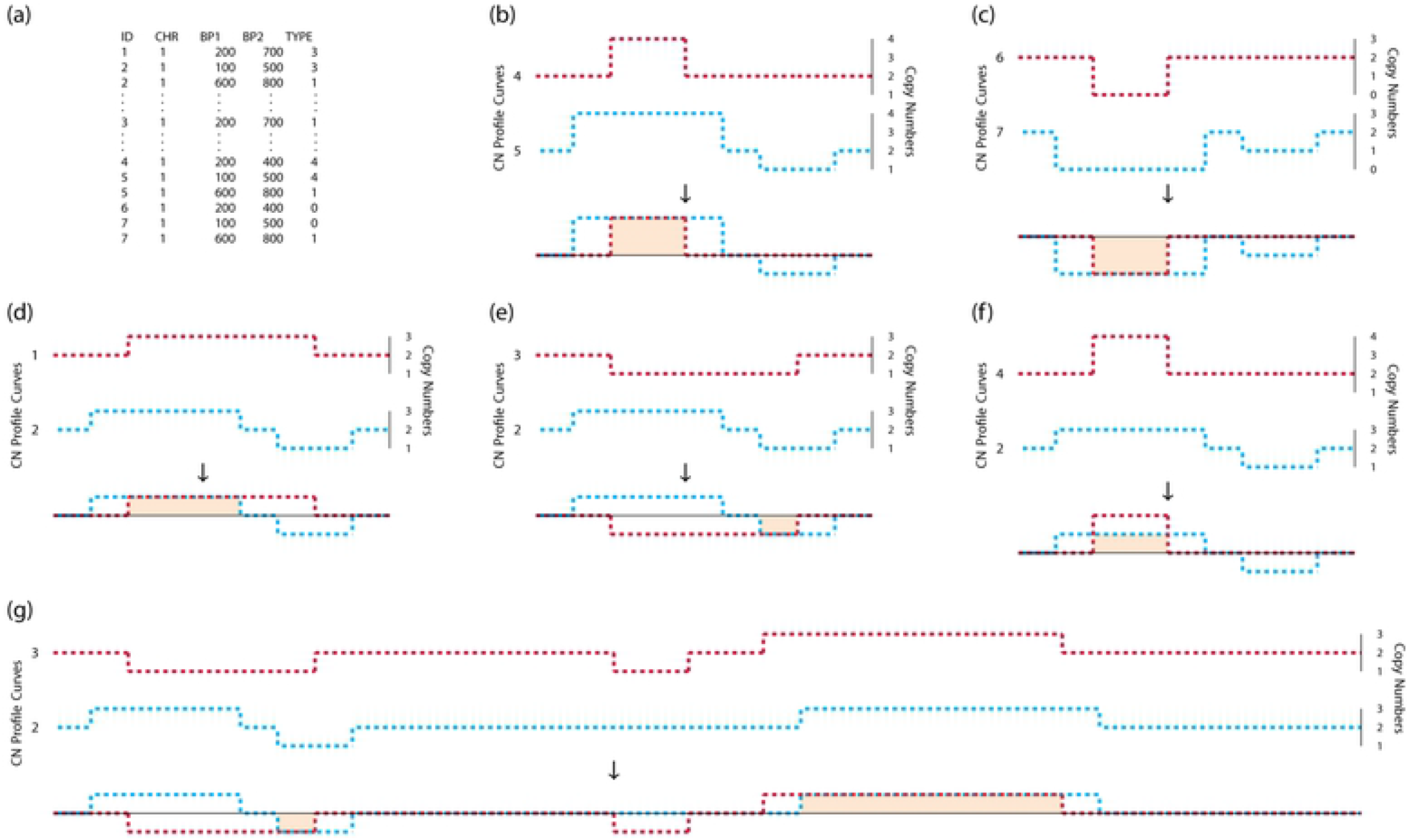
Diagram of copy number profile curves and common area under the curve. (a) Example of CNV data in standard PLINK format describing profiles of individuals in a small region of chromosome 1. (b)&(c) Copy number (CN) profile curves illustrating the cAUC between individuals with overlapping duplications of dosage 4 in (b) and individuals with overlapping deletions of dosage 0 in (c). (d)&(e) CN profile curves illustrating the cAUC between individuals with overlapping duplications of dosage 3 in (d) and individuals with overlapping deletions of dosage 1 in (e). (f) CN profile curves illustrating the cAUC between individuals with overlapping duplications of dosage 3 and 4. (g) CN profile curves which contain overlapping CNVs in multiple locations, so that the cAUC between the individuals is the sum of the two areas.

By superimposing two CN profile curves, we identify regions of overlapping CNVs of the same type (i.e., deletion or duplication) and propose to use the common area under the curve (cAUC) to quantify CNV similarity between two individuals. To implement the idea, first the raw dosage values in the CN profile curve are centered and scaled to obtain the duplication profile curve and deletion profile curve. The scaling and centering can be achieved by the dosage (DS) transform functions: *a^Dup^*(*DS*) = (*DS* − 2)^*d*^ for duplications and 0 otherwise, and *a^Del^*(*DS*) = (2 − *DS*)^*d*^ for deletions and 0 otherwise, where *d* is some pre-specified constant. Second, we superimpose the duplication profile curves of two individuals and note the overlapping regions where both curves are non-zero. Third, for each overlapping region, we multiply the minimum of the two respective transformed dosage values by the length of the overlap, and save this measure of “area of commonality”. Finally, we calculate the cAUC between two individuals as the sum of all such areas of commonality in their duplication profile curves plus the sum of all areas in their deletion profile curves. In the special case with *d* = 1 in the dosage transform functions *a^Dup^*(*DS*) and *a^Del^*(*DS*), the cAUCs between various pairs of individuals are illustrated in Fig 1. For individuals with overlapping CNVs of dosage 4 (for duplications; Fig 1 (b)) or dosage 0 (for deletions; Fig 1 (c)), the cAUC is the overlapping length times 2. For individuals with overlapping CNVs of dosage 3 (for duplications; Fig 1 (d)) or dosage 1 (for deletions; Fig 1 (e)), the cAUC is the overlapping length times 1. The cAUC between individuals with overlapping CNVs of the same type but different dosages (e.g., 3 versus 4), is the length of the overlap times 1 (Fig 1 (f)). If there are multiple overlaps in the individuals’ CN profile curves, the cAUC between two individuals is the sum of all areas of commonality (e.g., sum of shaded regions in Fig 1 (g)). The cAUC kernel measures similarity in both CNV length and dosage and hence characterizes the joint dosage and length effects. Using the semi-parametric kernel machine regression framework, CONCUR regresses the trait values on CNV effects captured by the cAUC kernel and evaluates the association between traits and CNV profiles via a score-based variance component test.

### Simulation design

The simulations were based on the pseudo-CNV data of 2000 individuals which is publicly available at https://www4.stat.ncsu.edu/~jytzeng/Software/CCRET/software_ccret.php. Autosome-wide pseudo-CNV data were simulated by mimicking the CNV profiles of unrelated individuals in the TwinGene study [8], and details are described in Tzeng et al. [6]. Briefly, the TwinGene study used a cross-sectional sampling design and included over 6,000 unrelated subjects born between 1911 and 1958 from the Swedish Twin Registry [9,10]. CNV calls were generated using Illumina OmniExpress beadchip for 72,881 SNP markers and using PennCNV (version June 2011) [11] as the CNV calling algorithm with recommended model parameters. From the full callset, high quality rare CNVs (frequency < 1% and size > 100kb) were extracted to form the simulation pool for the pseudo-CNV data. By mimicking the CNV profiles observed in a population dataset such as TwinGene, the pseudo-CNV data are appropriate for the simulation studies in this work. The pseudo-CNV data are stored in PLINK format indicating individual ID, CNV chromosome and starting and ending locations in base pairs (bp), and CNV dosage (e.g., 0, 1, 2, 3, etc.).

For the purpose of simulations we constructed “CNV segments” based on the pseudo-CNV profiles. The endpoints of the segments correspond to locations where a CNV in any one of the samples begins or ends, resulting in segments that contain either one or more intersecting CNVs. Within a segment, CNV dosage of an individual is a constant, and CNVs across individuals may have different dosages but share the same starting and ending positions. Note that different segments will naturally have different lengths. In the simulation study, we built design matrices **Z**^*Dup*^, **Z**^*Del*^, and **Z**^*Len*^ which codified CNV features by segment in the pseudo-CNV profile data. The dosage matrices took value 0 for those individuals without CNVs in the segment and were coded as 1 or 2 according to the number of additional or missing copies comprising the CNV. Length was the length of the CNV segment in kb for individuals with CNV events and was 0 for individuals without CNVs in the segment.

A case-control phenotype was generated from the logistic model

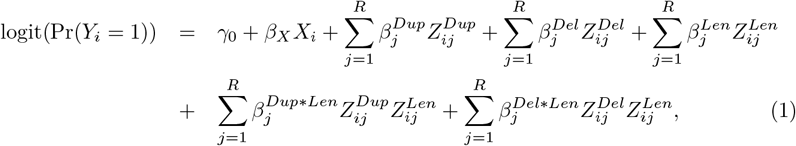

where *Z_ij_*^•^ is the (*i, j*) entry of matrix **Z**^•^, *i* = 1, ⋯, *N* indexes individuals, and *j* = 1, ⋯, *R* indicates CNV segment. A binary covariate *X_i_* was simulated from Bernoulli(0.5) for each individual. 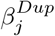 and 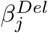 are the log-odds ratios of segment *j* for the presence of a CNV versus the absence. Likewise, 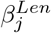 controls the effect of CNV length in segment *j*, and 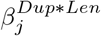 and 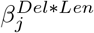 allow the effects of CNV length to differ by dosage. *β_j_*^•^ > 0 (or < 0) corresponds to a deleterious (or protective) CNV effect, and *β_j_*^•^ was set to 0 in non-causal segments. We set *β_X_* = log(1.1) and *γ*_0_ = −2, which corresponds to a disease rate of exp(−2) = 13.5% in baseline population. We also fixed 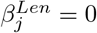 to reflect the observation that length tends to act like an effect modifier of dosage effects.

Among the CNV segments across the genome, we selected 200 segments to be causal, which consist of 100 causal “dup-segments” with at least one duplication and another 100 causal “del-segments” with at least one deletion. A causal dup-segment cannot be a causal del-segment. These causal segments were chosen as a random draw of 50 pairs of adjacent segments which both contained duplications, and another 50 pairs of adjacent segments which both contained deletions. This adjacent causal segment approach was designed to ensure that causal regions had more realistic lengths, since some segments were very short by chance.

We compare the performance of CONCUR with CCRET and CKAT. To implement CCRET, we used the functions from the CCRET package to convert the PLINK data to CCRET design matrices and computed the dosage kernel matrix. For CKAT, following Zhan et al. [7], we designated each chromosome as a CNVR and performed an association test for each chromosome. We reported the Bonferroni-corrected p-value for an overall association by multiplying the minimum p-value among the 22 association tests by 22. CNV lengths within each chromosome were scaled to be in [0,1] by dividing by the range of each chromosome, i.e., the maximal ending position minus the minimal starting position of observed CNVs on each chromosome. The Gaussian kernel scaling parameter was set to be 1.

We examined the methods’ performance under two signals: in Scenario I under a dosage×length signal and in Scenario II under a dosage-only signal. We chose these signals to roughly replicate the simulation settings applied to assess CKAT in [7] (dosage×length signal) and to assess CCRET in [6] (dosage signal). Under each scenario, we considered three sub-scenarios: (a) causal duplication effects only (referred to as Scenario I.a or II.a); (b) causal deletion effects only (referred to as Scenario I.b or II.b); and (c) both duplications and deletions to be causal (referred to as Scenario I.c and II.c). Within each sub-scenario, we varied the percentage of deleterious and protective effects by letting a percentage of the causal segments be deleterious or protective. We considered (1) 100% deleterious effects, (2) 50% deleterious and 50% protective, and (3) 10% deleterious and 90% protective. The choice of asymmetric heterogeneity settings was motivated by the rarity of 100% protective CNV effects in a genome-wide analysis, whereas 100% risk-associated effects are not uncommon. The power was evaluated in the range of odds ratios (exp(*β*)) 1.02-1.10 for Scenario I (dosage×length effects) and 1.1-1.9 for Scenario II (dosage effects). Power estimates are reported for a range of effect sizes such that the power ranges roughly from 0.2 to 0.8.

We implemented case-control sampling to obtain 2000 cases and 2000 controls for each simulation replication. Type I error rates were evaluated based on 5000 replications, and power was estimated based on 300 replications at each effect size. For all methods (i.e., CONCUR, CCRET and CKAT), we adjusted for a simulated binary covariate as a fixed effect in the kernel machine regression. We employed the small-sample variance components test of Chen et al. [12] and obtained p-values using Davies’ method [13] as implemented in the CKAT R package.

### Simulation Results

The type I error rates of the three tests were examined at nominal levels of 0.01, 0.05, and 0.1 (Table 1). All methods had type I error rates roughly around the nominal level.

**Table 1.** Type I error rates. Type I error rates of three CNV tests evaluated based on 5000 replications.

| Nominal level | CONCUR | CCRET | CKAT |
| --- | --- | --- | --- |
| 0.01 | 0.010 | 0.008 | 0.009 |
| 0.05 | 0.045 | 0.047 | 0.049 |
| 0.10 | 0.096 | 0.093 | 0.092 |

#### Scenario I: Causal Dosage×Length Effects

Scenario I.a (I.b) considers dosage-length interactions only from causal duplication (deletion) segments, and includes three settings of mixed deleterious and protective effects which are labeled as (D,P)=(100,0), (50,50) and (10,90); (D,P) indicates the proportion of deleterious (D) and protective (P) segments among all causal segments. The results are displayed in Fig 2, with the top row showing power under causal duplication effects and bottom row under causal deletion effects. The CONCUR method has the best or comparable power with the second best method (CCRET) across different settings of deleterious-protective effects. Both CONCUR and CKAT are designed to detect dosage×length signals, but CKAT struggled to pick up this signal perhaps due to applying the method to very large CNVRs (chromosomes) as well as the multiple testing penalty. We also observed a difference in relative performance in the (D,P)=(50,50) setting between I.a (causal duplications) and I.b (causal deletions). This is not unexpected because in the simulated data, there are differences in the features of duplication and deletion events. The proportion of the causal deletion sites out of all deletions was 9.5%, and is 6.9% for duplications. In addition, the 100 causal duplication segments had higher median and mean length compared to the 100 causal deletion segments (median 75kb vs. 32kb; mean 81kb vs. 64kb).

**Fig 2.**
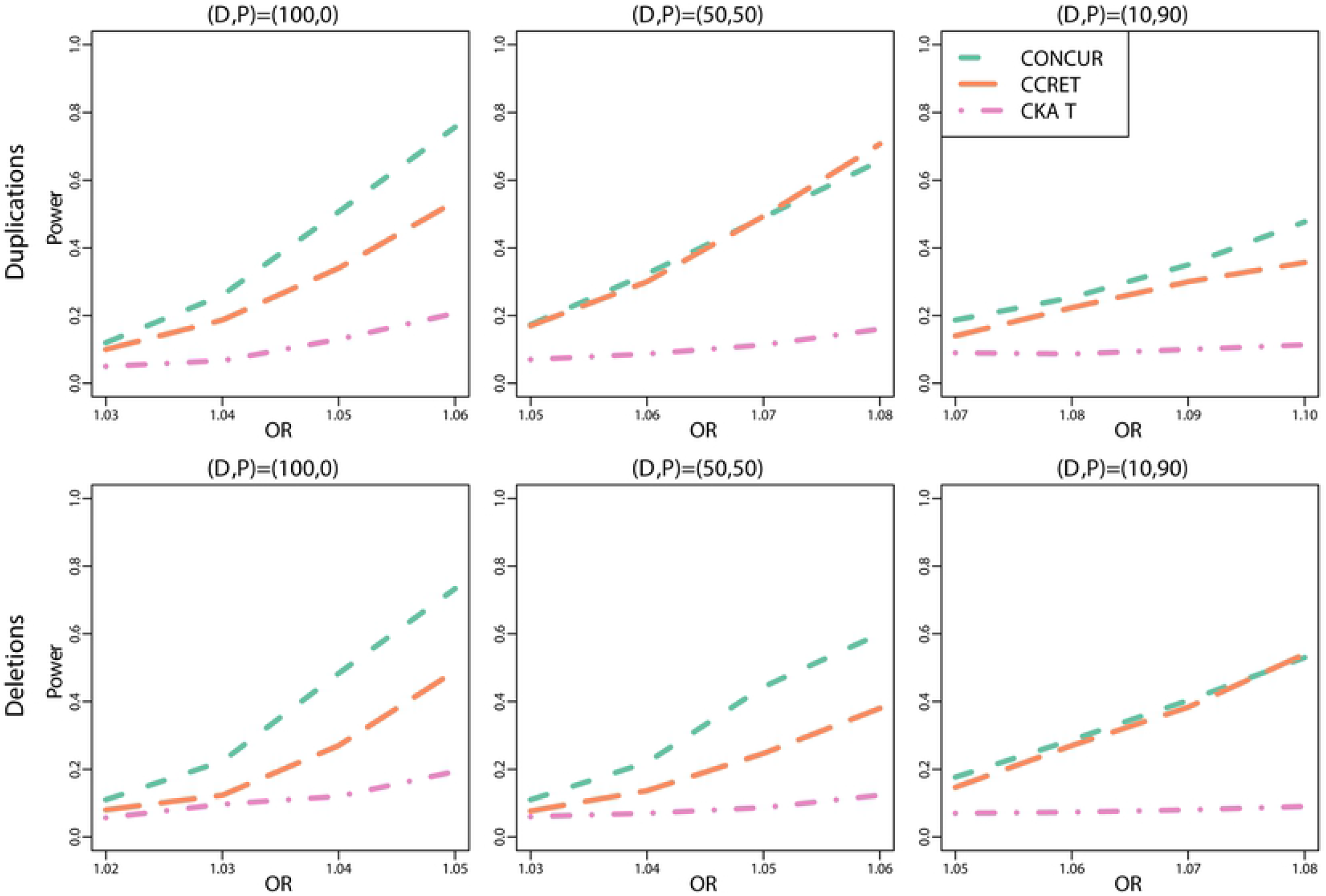
Power comparison between CONCUR, CCRET, and CKAT under Scenario I (causal dosage×length effects). The top panel shows results from Scenario I.a (causal duplication effects) and the bottom panel from Scenario I.b (causal deletion effects). In each sub-scenario, three different proportions of deleterious vs. protective effects are considered as indicated by (D,P), with D representing the proportions of deleterious segments and P the protective segments among causal segments.

Scenario I.c considers dosage-length interactions from both duplications and deletions and includes four settings of mixed deleterious and protective effects. (Fig 3) These settings are denoted as (D_Dup_,P_Dup_,D_Del_,P_Del_)=(100,0,100,0), (50,50,50,50), (90,10,10,90), and (10,90,90,10), where D_Dup_ and P_Dup_ respectively are the proportions of deleterious and protective segments among causal duplication segments, and D_Del_ and P_Del_ are defined similarly for causal deletion segments. These settings allow the assessment of the method performance under multiple sources of effect heterogeneity, including between-locus heterogeneity due to the mixture of deleterious and protective segments, between-locus heterogeneity due to duplication and deletion causal segments, and within-locus heterogeneity due to duplications and deletions with a segment having opposite effects. We observed that CONCUR has the best power among the three tests across different settings, followed by CCRET and then by CKAT.

**Fig 3.**
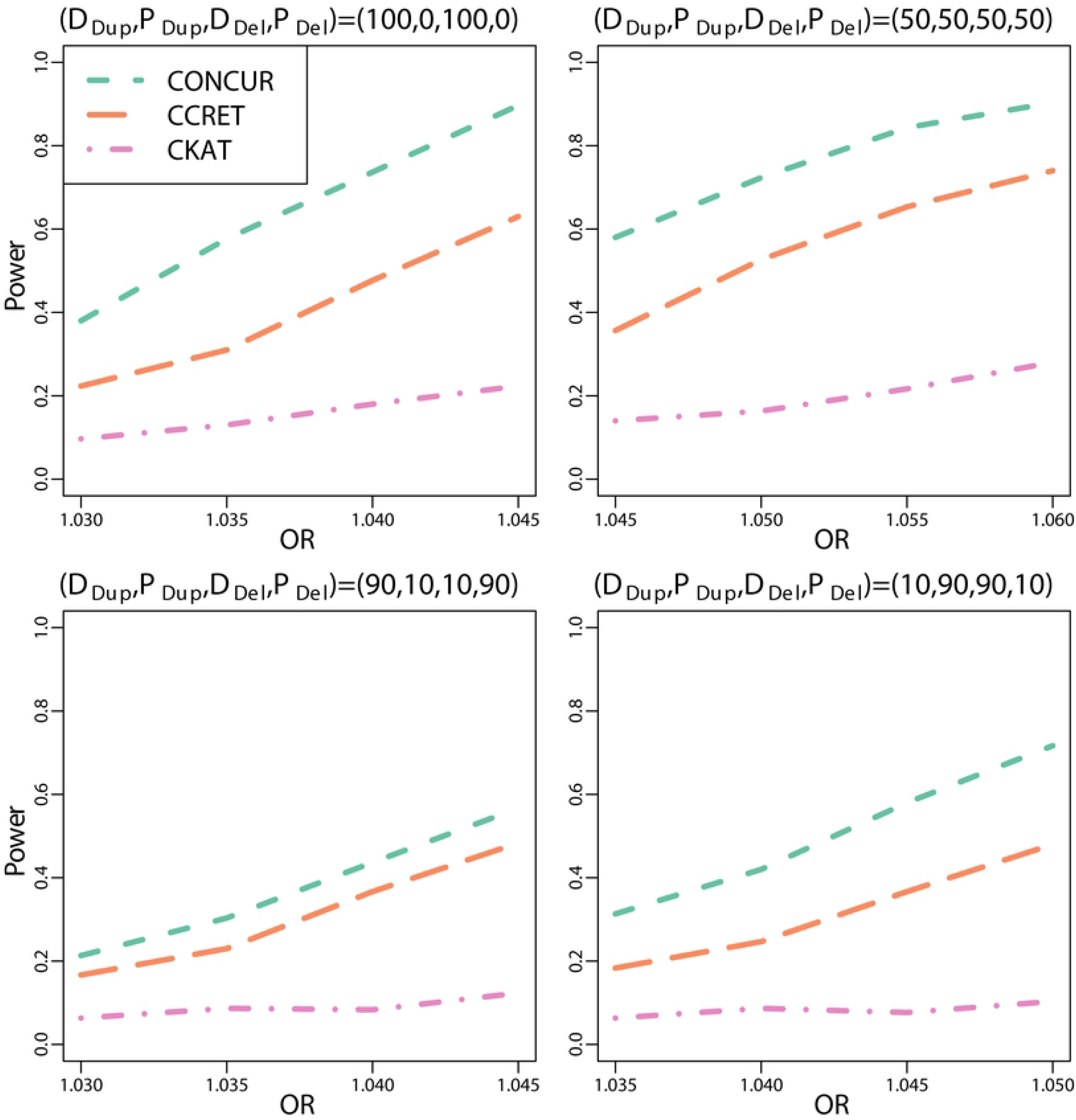
Power comparison between CONCUR, CCRET, and CKAT for Simulation I.c (causal dosage×length effects from both duplications and deletions). Four different proportions of deleterious vs. protective effects are considered as indicated by (D_Dup_,P_Dup_,D_Del_,P_Del_) with D_Dup_ and P_Dup_ reflecting the proportions of deleterious and protective segments among causal duplication segments, and with D_Del_ and P_Del_ defined similarly for causal deletion segments.

#### Scenario II. Causal Dosage Effects

Scenario II.a (II.b) considers dosage effects from causal duplication (deletion) segments, and includes three settings of mixed deleterious and protective effects, i.e., (D,P)=(100,0), (50,50) and (10,90). The results are shown in Fig 4. As expected, the dosage-based CCRET kernel performs the best, with CONCUR following CCRET or having comparable power. Similar results are observed under Scenario II.c (Fig 5), where causal dosage effects are from both duplications and deletions and four varying mixtures of deleterious and protective effects are considered.

**Fig 4.**
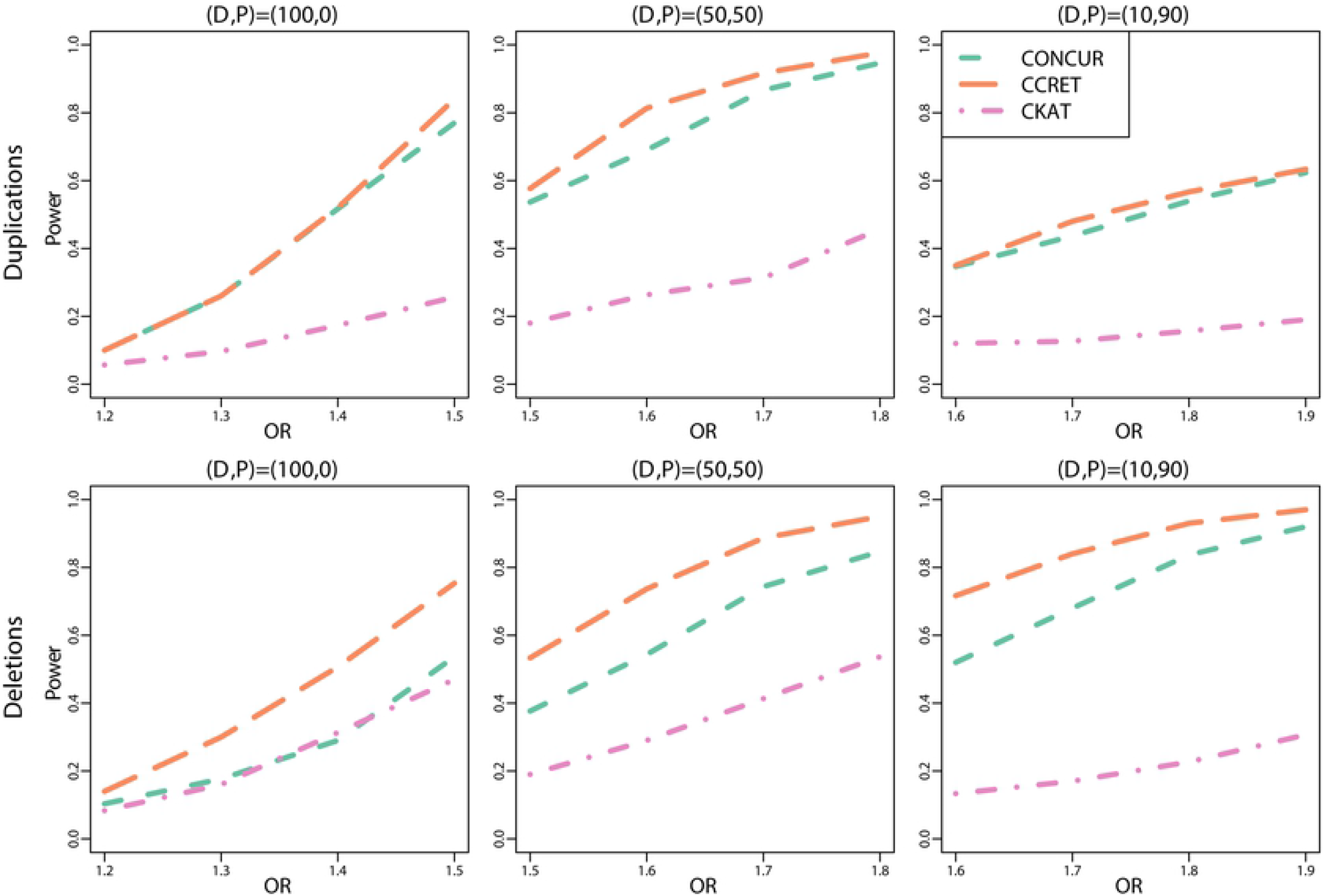
Power comparison between CONCUR, CCRET, and CKAT under Scenario II (causal dosage effects). The top panel shows results from Scenario II.a (causal duplication effects) and the bottom panel from Scenario II.b (causal deletion effects). In each sub-scenario, three different proportions of deleterious vs. protective effects are considered as indicated by (D,P), with D representing the proportions of deleterious segments and P the protective segments among causal segments.

**Fig 5.**
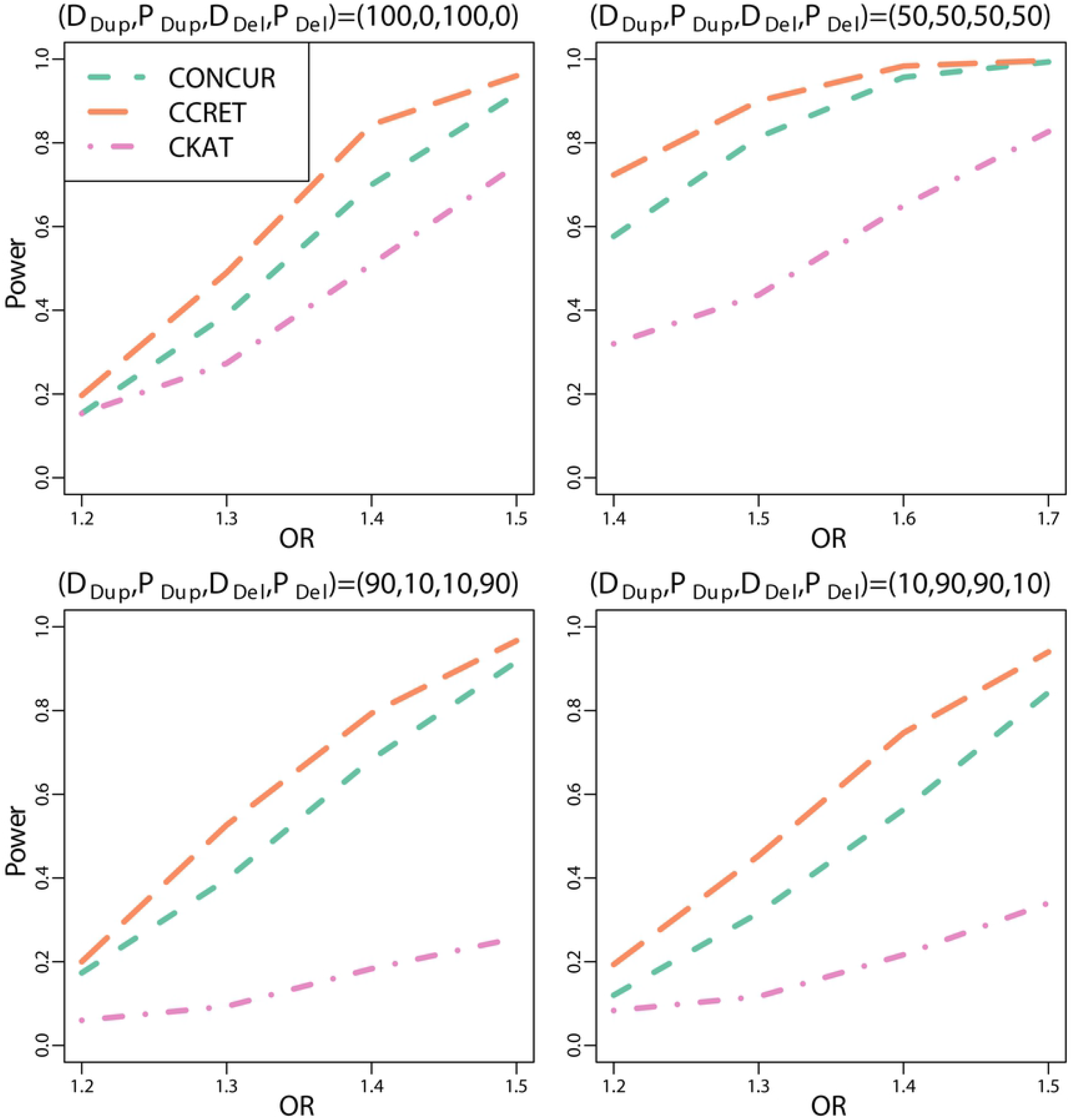
Power comparison between CONCUR, CCRET, and CKAT for Simulation II.c (causal dosage effects from both duplications and deletions). Four different proportions of deleterious vs. protective effects are considered as indicated by (D_Dup_,P_Dup_,D_Del_,P_Del_) with D_Dup_ and P_Dup_ reflecting the proportions of deleterious and protective segments among causal duplication segments, and with D_Del_ and P_Del_ defined similarly for causal deletion segments.

### Real data application

In real data applications, we first, as a proof of concept, applied the proposed CONCUR test on a previously analyzed CNV dataset from the Swedish Schizophrenia Study. We next conducted a CNV-triglyceride (TG) association analysis using CONCUR on data from the Taiwan Biobank.

### CNV analysis on schizophrenia in the Swedish Schizophrenia Study

We conducted pathway-based CNV analysis on data from the Swedish Schizophrenia Study [14]. The Swedish Schizophrenia Study used a case-control sampling design. Genotyping was done in six batches using Affymetrix 5.0 (3.9% of the subjects), Affymetrix 6.0 (38.6%), and Illumina OmniExpress (57.4%). PennCNV [11] was used to generate CNV calls. After quality control, we obtained a high quality rare CNV (frequency < 1% and size > 100kb) dataset in 8,547 subjects (3,637 cases and 4,820 controls) [15]. All procedures were approved by ethical committees at the Karolinska Institutet (Dnr No. 04/-449/4 and No. 2015/2081-31/2) and University of North Carolina (No. 04-1465 and No. 18-1938). All subjects provided written informed consent (or legal guardian consent and subject assent). Previous analyses of this data [15] indicated significant associations of large rare CNVs with schizophrenia risk for both genome-wide dosage effects and gene intersecting effects of selected gene sets.

To evaluate the practical utility of the three kernel-based tests, we performed analysis on the gene sets previously examined in [6], excluding the PSD pathway as it overlaps the other three PSD-related pathways considered. In the eight gene sets, large (> 500kb) rare CNVs were found to be associated with schizophrenia by Szatkiewicz et al. [15], and these associations were corroborated by Tzeng et al. [6] in a gene-interruption analysis with CNVs > 100kb. In each pathway analysis, we performed association tests for joint dosage and length effects of rare CNVs > 100kb, using a fixed effect term to adjust for batch effects. CONCUR and CKAT kernels were constructed from the raw PLINK data and the CCRET dosage kernel was created using the functions available on the CCRET website. For CKAT, we used pathways as the CNVR unit instead of chromosomes because there were multiple chromosomes with only one gene. The results were evaluated against a Bonferroni-adjusted threshold of 0.05/8 = 0.00625.

CONCUR found significant associations in all pathways, while CCRET and CKAT had alternating significance in some of the pathways (Table 2). In the FMRP pathway, all three tests were significant, and in the remaining seven gene sets, one or both of CCRET and CKAT were significant or near significant. The analyses suggest significant CNV effects from dosage and/or length affecting schizophrenia risk, and the relative performance of these methods suggest some implications about the underlying effect patterns. CKAT, which is more sensitive to dosage-length interactive effects, found slightly more and different significant associations compared to CCRET, which is more sensitive to dosage effects, while CONCUR appeared to be more encompassing. We also observed stronger power of CKAT in the analysis here compared to the power observed in the simulation studies, which may partially be due to the lack of multiple testing penalty here.

**Table 2.** Association test results for the effects of CNVs with > 100kb in length on schizophrenia risk in the Swedish Schizophrenia Study. Pathways are ordered by the number of tests that found significance (3 tests, 2 tests, 1 test) and then by pathway name. Significant p-values (at threshold 0.05/8=0.00625) are shown in bold.

| Gene-set Name | Gene-sets |  |  | P-values |  |  |
| --- | --- | --- | --- | --- | --- | --- |
|  | # Genes | # Genes Interrupted<br>in Cases | # Genes Interrupted<br>in Controls | *CONCUR* | *CCRET* | *CKAT* |
| FMRP targets (Darnell et al. [16]) | 810 | 149 | 152 | **2.29E-05** | **0.00044** | **0.00026** |
| PSD/PSD-95 (Kirov et al. [17]) | 65 | 13 | 10 | **0.00052** | **0.00144** | 0.00903 |
| Synaptic Proteome (G2Cdb) | 1023 | 121 | 106 | **0.00067** | **0.00010** | 0.00736 |
| Cytoplasm (Kirov et al. [17]) | 266 | 28 | 32 | **0.00124** | 0.01408 | **0.00030** |
| Mental Retardation | 503 | 67 | 63 | **0.00164** | 0.10200 | **0.00350** |
| PSD/mGluR5 (Kirov et al. [17]) | 38 | 4 | 7 | **0.00040** | 0.10540 | **0.00129** |
| PSD/NMDAR (Kirov et al. [17]) | 61 | 12 | 12 | **0.00102** | 0.00922 | **0.00046** |
| Synaptic genes (Ruano et al. [18]) | 718 | 154 | 164 | **5.45E-06** | 0.02005 | 0.00766 |

### CNV analysis on triglycerides in the Taiwan Biobank

We applied the proposed CONCUR test to the Taiwan Biobank (TWB) data https://www.twbiobank.org.tw/new_web/ and conducted CNV association analysis with triglyceride (TG) levels on lipid-related pathways. The nationwide biobank project was initiated in 2012 and has recruited more than 15,995 individuals. The study has been approved by the ethical committee at Taichung Veterans General Hospital (IRB TCVGH No. CE16270B-2). The consent was not obtained because the data were analyzed anonymously. Peripheral blood specimens were extracted from healthy donors and genotyped using the Affymetrix Genomewide Axiom TWB array, which was designed specifically for a Taiwanese population. The TWB array contains 653,291 SNPs and was used to generate calls for genome-wide CNVs in the following process. First, Affymetrix Power Tools version 1.18.0 was used to produce a summary file of the intensity values of all probes, and the file was input into the Partek Genomic Suite version 6.6 to call CNVs based on the following criteria: at least 35 consecutive SNP markers, p-values of different CN values between two consecutive segments < 0.001, and signal-to-noise ratio (SNR) ≥ 0.3. A duplication was called if its copy number was > 2.3, whereas a deletion was called if its copy number was < 1.7. Several previous studies [19] [20] have demonstrated appropriate CNV calls with these parameters. After quality control, we obtained CNV data in 14,595 unrelated individuals. Our CNV association analyses focused on a subset of 11,664 individuals who had non-missing TG levels.

We referenced the Kyoto Encyclopedia of Genes and Genomes (KEGG) pathway database [21] to identify lipid-related pathways. Among the 17 pathways related to “Lipid metabolism”, 15 pathways included genes intersected by the TWB CNV data and were selected. For each pathway we performed the CONCUR test, CCRET, and CKAT. We adjusted for sex, age, BMI, and the top 10 principal components representing the population structure as covariates with fixed effects. As before, CKAT was performed with each pathway comprising a single CNVR. We compared the test results to a Bonferroni threshold of 0.05/15 = 0.00333.

Out of the 15 pathways, ten pathways were identified as significantly associated with TG by CONCUR, nine pathways by CKAT, and one pathway by CCRET (Table 3). There were a total of 12 pathways found significant by one or more methods, among which one pathway, hsa00120 (primary bile acid biosynthesis), was significant for all methods. Compared to the Swedish Schizophrenia Study analysis, CCRET suffered from lower power and CKAT showed greater power, while the performance of CONCUR was relatively stable. The power loss in CCRET might be due to more dominant length or dosage×length signals and perhaps also a consequence of the stricter significance threshold here. CKAT demonstrated much better power than in the simulation study, which is likely attributable to the treatment of each pathway as a CNVR and hence the absence of multiple testing adjustment needed for multiple CNVRs. However, although CONCUR and CKAT were significant in a roughly equal number of pathways (ten versus nine, respectively), the CONCUR p-values tended to be much smaller than the CKAT p-values. To illustrate, if a more stringent significance threshold was adopted to adjust for the total of 45 tests (15 pathways × 3 methods) at a Bonferroni threshold of 0.05/45=0.0011, then CONCUR would maintain significance in seven pathways while no CKAT p-value would meet the threshold. This behavior somewhat echoes the performance of CKAT in the simulation study.

**Table 3.** Association test results for the effects of CNVs on triglyceride levels in the Taiwan Biobank. Pathways are ordered by the number of tests that found significance (3 tests, 2 tests, 1 test, and no tests) and then by pathway name. Significant p-values (at threshold of 0.05/15 = 0.00333) are shown in bold.

| Gene-sets |  |  |  | P-values |  |  |
| --- | --- | --- | --- | --- | --- | --- |
| Gene-set Names | # Genes | # Genes Interrupted |  | *CONCUR* | *CCRET* | *CKAT* |
| hsa00120<br>(Primary acid bile biosynthesis) | 17 | 17 |  | **0.00019** | **0.00314** | **0.00274** |
| hsa00061<br>(Fatty acid biosynthesis) | 13 | 12 |  | **0.00171** | 0.01187 | **0.00197** |
| hsa00140<br>(Steroid hormone biosynthesis) | 60 | 58 |  | **0.00030** | 0.00623 | **0.00159** |
| hsa00564<br>(Glycerophospholipid metabolism) | 97 | 86 |  | **0.00322** | 0.00398 | **0.00209** |
| hsa00590<br>(Arachnidonic acid metabolism) | 63 | 62 |  | **0.00212** | 0.00883 | **0.00211** |
| hsa00591<br>(Linoleic acid metabolism) | 29 | 29 |  | **0.00080** | 0.01799 | **0.00291** |
| hsa01040<br>(Biosynthesis of unsaturated fatty acids) | 27 | 23 |  | **0.00012** | 0.00394 | **0.00158** |
| hsa00062<br>(Fatty acid biosynthesis) | 30 | 26 |  | **0.00031** | 0.01591 | 0.00508 |
| hsa00072<br>(Synthesis and degradation of ketone bodies) | 10 | 10 |  | **0.00008** | 0.00459 | 0.00383 |
| hsa00561<br>(Glycerolipid metabolism) | 61 | 50 |  | 0.00430 | 0.00494 | **0.00198** |
| hsa00565<br>(Ether lipid metabolism) | 47 | 43 |  | **0.00018** | 0.00859 | 0.00439 |
| hsa00592<br>(alpha-Linolenic acid metabolism) | 25 | 25 |  | 0.00581 | 0.00927 | **0.00273** |
| hsa00071<br>(Fatty acid degradation) | 44 | 43 |  | 0.00406 | 0.01088 | 0.00631 |
| hsa00100<br>(Steroid biosynthesis) | 19 | 16 |  | 0.01641 | 0.00618 | 0.00906 |
| hsa00600<br>(Sphingolipid metabolism) | 47 | 43 |  | 0.00382 | 0.00512 | 0.00789 |

The relative performance of CONCUR, CKAT and CCRET seems to suggest that CNV length or dosage×length effects dominate in the 12 significant pathways. To illustrate possible CONCUR post hoc analyses so to probe the potential sources of the pathway-level signal, we looked more closely at one pathway, hsa01040 (biosynthesis of unsaturated fatty acids), for which both CONCUR and CKAT were significant while CCRET was borderline significant. Previous studies have reported that monounsaturated fat acids or polyunsaturated fatty acids can effect TG levels [22,23]. Given the major function of the genes in hsa01040 (i.e., the biosynthesis of unsaturated fatty acids), it is not unexpected that CNVs in these genes were significantly associated with TG levels. We calculated summary statistics describing CNV length and dosage in hsa01040 for individuals with different levels of TG. Based on the TG quantiles from the sample data, we classified individuals as having high TG (>75th percentile [>140 mmHg]), medium TG (25th–75th percentile [68-140 mmHg]) and low TG (<25th percentile [<68 mmHg]). We applied ANOVA to detect differences in CNV length and in dosage characteristics, and applied chi-squared tests to assess differences in the proportion of individuals with CNVs across TG levels. In addition, we examined CNV features in all CNVs together and in duplications and deletions separately.

**Table 4.** Descriptive statistics for hsa01040 pathway. TG values are classified as Low (<the 25th percentile [<68 mmHg]; n=2,931), Medium (the middle 50% [68 - 140 mmHg]; n=5,844), and High (>the 75th percentile [>140 mmHg]; n=2,889). The percent of individuals with CNVs is with respect to the total number of individuals in each TG category. The mean number of CNVs per individual and mean total length of CNVs (bp) per individual are reported, as well as the mean lengths (bp) and mean dosage per CNV. “Promising” associations with TG are marked with ⋆⋆ to indicate p-value< 0.01 and with ⋆ to indicate p-value< 0.05.

| CNV Type | TG Level | Pct<br>Individuals<br>with CNV | Mean #<br>CNVs per<br>Individual | # Genes<br>Interrupted | Mean Total<br>CNV Length<br>per Individual (bp) | Mean CNV<br>Length (bp) | Mean CNV<br>Dosage |
| --- | --- | --- | --- | --- | --- | --- | --- |
| All | Low | 6.18% | 3.33 | 23 | 25143.71 | 2433.58 | 1.63 |
|  | Medium | 6.07% | 3.52 | 23 | 24447.30 | 2473.48 | 1.63 |
|  | High | 7.17% | 3.48 | 23 | 31091.65 | 2471.43 | 1.64 |
| Deletion | Low | 2.8% | 5.84 | 16 | 29630.62 | 2590.55★ | 1.41 |
|  | Medium | 2.74% | 6.24 | 17 | 28107.36 | 2593.05★ | 1.40 |
|  | High | 3.32% | 5.79 | 16 | 32039.63 | 2067.96★ | 1.39 |
| Duplication | Low | 3.62% | 1.17 | 20 | 7811.20 | 1827.24★★ | 2.50★ |
|  | Medium | 3.54% | 1.22 | 23 | 10009.61 | 2001.81★★ | 2.52★ |
|  | High | 4.15% | 1.38 | 22 | 27897.23 | 3831.02★★ | 2.49★ |

Taking p-values < 0.05 as a suggestive “promising” association with TG, we did not observe any CNV associations when all CNVs were analyzed together, but for duplications only, there were promising differences in CNV length (p-value=0.0063) and weaker differences in dosage (p-value=0.0255) across TG levels. There were also some weak significance in CNV length for deletions (p-value=0.0423). We were cautious to not over-interpret these “promising” associations since this stratified analysis reflected only marginal associations of a CNV feature, and the tests did not account for the effect heterogeneity that motivates the application of kernel-based methods. We also proceeded with testing using CONCUR on duplications and deletions separately, and found a very significant association with TG in duplications (p-value < 1 × 10^−8^) and a weaker signal in deletions (p-value=0.0313).

To further explore the signal from duplications, we visualized CNVs in the 23 genes in hsa01040 (Fig 6). Fig 6 displays duplications and deletions in the CNV profiles of individuals categorized by their TG level (low, medium, and high), with profiles clustered so that shared patterns across profiles become apparent. For exploration purposes, we applied CONCUR to duplications in each gene and found that several genes had strong association p-values (i.e., < 10^−4^), *BAAT, ELOVL4, ELOVL6, ELOVL5, HSD17B4*, and *SCD5* (S1 Table). Notably, *BAAT* is an amino acid N-acyltransferase for bile acid. Previous studies have demonstrated that bile acids are important regulators for TG level through crosstalk with farnesoid X receptor (FXR) [24,25]. Since conversion of cholesterol to bile acid is an essential step in preventing the accumulation of TG, copy number duplications in *BAAT* may directly affect TG levels in the blood. Three *ELO* genes had significant CNV associations. Since the major functions of these genes focus on the elongation of fatty acids, CNV events in these genes are likely to affect the production and metabolism of TG. For example, one study showed that hepatic steatosis was observed in *ELOVL5*-knockout mice due to the activation of SREBP-1c and its target genes [30]. *HSD17B4* is a dehydrogenase, which is able to inhibit the production of DHEA [26]. A previous study showed that TG levels were inversely correlated to DHEA levels in men with type 2 diabetes [27], suggesting a potential link between CNVs in *HSD17B4* and TG levels. *SCD5* serves as a critical enzyme providing a double bond to construct complex lipid molecules such as TG [28,29], and thus dysregulation of *SCD5* expression may impact TG levels. Further analyses are required to formally localize the sources of the CNV association signal in this pathway and others, but this exploratory analysis nonetheless serves to enrich our understanding of the association in pathway hsa01040 through examination of CNV-level and gene-level features.

**Fig 6.**
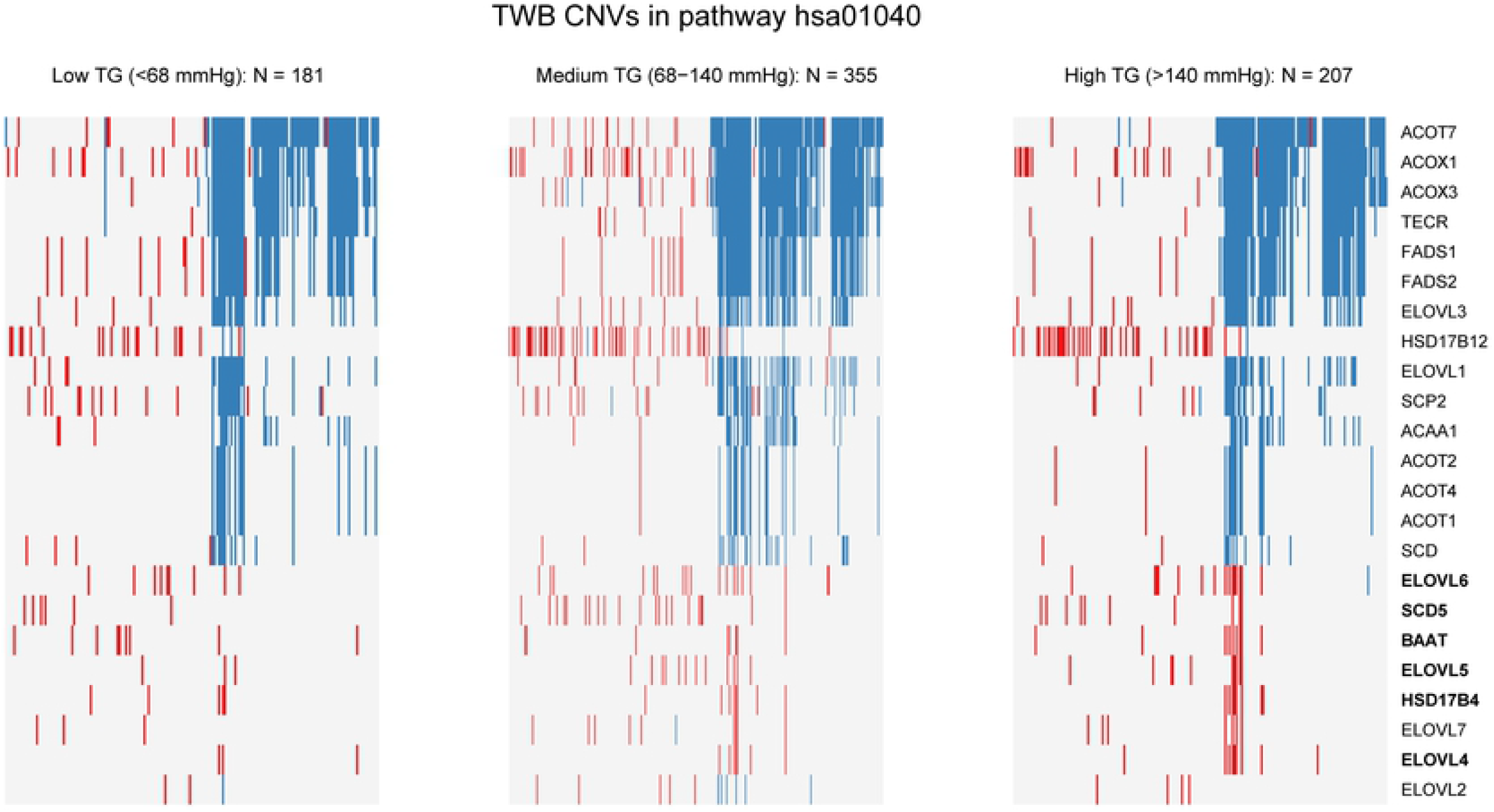
Visualization of CNV activity in pathway hsa01040 by level of triglycerides (TG). CNV activity in genes in hsa01040 is shown by level of TG (Low, Medium, and High), with duplications in red and deletions in blue. Columns represent clustered individuals, and genes shown here are the 23 genes in the pathway that contain CNVs, ordered by the number of CNVs contained therein.

## Discussion

We introduce CONCUR to leverage the strength of kernel-based methods to access the collective effects of rare CNVs on disease risk and incorporate several desired features. First, CONCUR permits the quantification of CNV similarity in an CNVR-free manner, avoiding the need of arbitrarily defining CNVRs as in current practice. Second, CONCUR incorporates both length and dosage information via the cAUC kernel, and is capable of detecting dosage, length and length-dosage interaction effects. Third, as the technology for detecting smaller CNVs improves, we expect to observe more length variation in CNVs and an increasing need to accommodate length effects in CNV association studies. However, there exist shortcomings in the standard kernel choices for handling CNV length. For example, a linear (or polynomial) kernel, which scores length similarity in a multiplicative fashion, cannot always reflect the true level of length similarity between an individual pair, e.g., a pair of CNVs of length 20kb would be equally similar to two CNVs with lengths 1kb and 400kb (as 20×20 = 1×400). The alternative, e.g., Gaussian kernel as in CKAT, would still require a pre-specified scaling factor. CONCUR addresses these issues by using the common AUC of the CN profile curves of an individual pair and quantifies CNV similarity in dosage and length simultaneously. Finally, unlike current kernel methods, which require discretized copy numbers, CONCUR is directly applicable to continuous and discrete copy numbers. We provide the R functions that perform the CONCUR test at https://www4.stat.ncsu.edu/~sthollow/JYT/CONCUR/.

CONCUR shares some philosophy with several CNV analysis strategies in the literature. For example, Aguirre et al. [31] characterized the copy number changes in the pancreatic adenocarcinoma genome by detecting the minimum common regions (MCR) of recurrent copy number changes across tumor samples and using MCRs to prioritize genes that might be involved in pancreatic carcinogenesis. Harada et al. [32] also examined the minimal overlapping/common regions of frequent CNV activities among pancreatic cancer samples and among normal samples to identify candidate regions that might contain critical oncogenes or tumor suppressor genes. Furthermore, Mei et al. [33] proposed algorithms for identifying common CNV regions across individuals of homogeneous phenotypes for downstream association analysis. Built on similar concepts to these “common regions”, CONCUR quantifies CNV similarity between sample pairs based on the “size” of the common regions as reflected in congruent location and dosage, and provides an association test to evaluate dosage and length effects.

In the analyses performed in this study, we calculated the cAUC using CNV dosage values transformed by the functions *a^Dup^*(*DS*) = (*DS* − 2) for duplications and 0 otherwise, and *a^Del^*(*DS*) = (2 − *DS*) for deletions and 0 otherwise. That is, we used copy number 2 as a reference value, and defined CNV similarity as the overlapping CNV length scaled linearly according to the magnitude of dosage deviation from the reference value. However, CONCUR can be flexibly extended to accommodate other schemes of quantifying common area by adopting different *a*(·) functions in the calculation of the cAUC. For example, instead of a linear scaling with *a*^•^(*DS*) = |*DS* − 2|), one may consider a non-linear scaling by setting *a*^•^(*DS*) = |*DS* − 2|^*d*^, with *d* < 1 deflating and *d* > 1 enhancing the contributions of CNVs of more extreme gains/losses. Additionally, one can impose reference values other than 2, such as using 2.3 for duplications (e.g., by setting *a^Dup^*(*DS*) = (*DS* − 2.3) for duplications and 0 otherwise), and using 1.7 for deletions (e.g., by setting *a^Del^*(*DS*) = (1.7 − *DS*) for deletions and 0 otherwise). Finally, overlapping area may be further weighted by inverse frequencies when needed, to augment the contribution of overlap in regions of rare CNV activity or of CNVs with uncommon dosage.

## Materials and methods

### CONCUR method

For individual *i, i* = 1, ⋯, *n*, denote *Y_i_* the phenotype of individual *i*. Codify the CNV information in matrix *Z_i_* with dimension *P_i_* × 4 as in the standard PLINK format of CNV data, where *P_i_* is the number of CNVs that individual *i* has, and each row of *Z_i_* records four features of CNV *p, p* = 1, ⋯, *P_i_*: dosage (denoted as *DS_p_*), chromosome (denoted *CHR_p_*), start location (denoted as *BP*1_*p*_), and end location (denoted as *BP*2_*p*_). The dosage *DS_p_* can be integer or continuous values. Finally let *X_i_* = (*X*_*i*1_, ⋯, *X_ir_*)^*T*^ be the *r* covariates. Under the kernel machine regression framework, we model the association between phenotypes and CNVs as follows

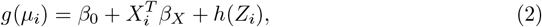

where *μ_i_* = *E*(*Y_i_*|*X_i_, Z_i_*), *g*(·) is the canonical link, and *h*(*Z_i_*) is an unknown smooth function of the variant features characterized by a kernel function *K*(·, ·). For continuous responses, *g*(*μ_i_*) = *μ_i_*; for binary responses, *g*(*μ_i_*) = log[(*μ_i_*/(1 − *μ_i_*)].

#### Profile curves

The proposed cAUC kernel is built on the concept of a CN profile curve. For a given chromosome *k* = 1, 2, ⋯, 22 and individual *i* = 1, 2, ⋯, *n*, we conceptualize a function 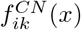 which returns the copy number of a CNV if *x* falls in a CNV and returns 2 (i.e., no CNV events) otherwise, e.g., examples shown in Fig 1. Given the CN profile curve, we further define two curves called the duplication profile curve and deletion profile curve, which recenter and rescale the CN values in CN profile curves through the “dosage transform functions” as described below, and allow us to compute cAUC similarity from duplications and from deletions in a more flexible manner.

We further use *q* = 1, ⋯, *P_ik_* to index the CNV features (*DS_q_, BP*1_*q*_, *BP*2_*q*_) occurring on chromosome *k* of individual *i* for *k* = 1, ⋯, 22. Then we construct duplication and deletion profile curves respectively describing duplications and deletions on chromosome *k* for individual *i* as follows:

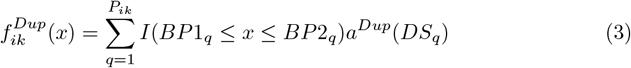

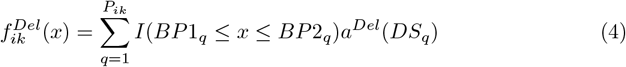

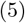

where *x* is a location on the genome on the same scale as *BP*1_*q*_ and *BP*2_*q*_; *I* is the indicator function such that *I*(·) = 1 if the condition contained within is satisfied and equals 0 if otherwise; and *a*^•^(*DS*) is a dosage transform function which determines the reference copy number value and controls how different copy number values contribute more or less to similarity in profiles. If an individual has no CNVs in chromosome *k*, then their duplication and deletion profile curves are identically equal to zero, i.e., 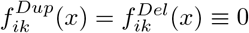 for all *x*. Although not explicitly shown, 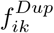 and 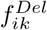 are functions of *Z_i_*, as the information of *DS_q_, BP*1_*q*_, *BP*2_*q*_ and chromosome *k* for subject *i* is obtained from *Z_i_*.

In this study, we designated *a^Dup^*(*DS_q_*) = (*DS_q_* − 2) if *DS_q_* is from a duplication and 0 otherwise and *a^Del^*(*DS_q_*) = (2 − *DS_q_*) if *DS_q_* is from a deletion and 0 otherwise. That is, for a given chromosome *k* and individual *i*, the function 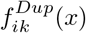 equals the magnitude of the duplication (i.e., number of additional copies compared to the reference copy number 2) for *x* inside a duplication and equals 0 otherwise, with analogous logic for 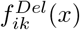. Other options of the dosage transform functions are described in the discussion section.

#### cAUC kernel

We propose to quantify the similarity between individuals *i* and *j* by comparing 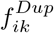 vs. 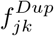 and 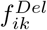 vs. 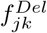 over chromosomes *k* =1, ⋯, 22 using the following kernel function

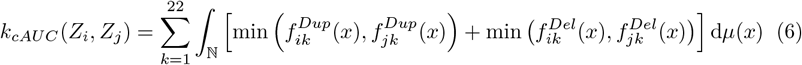

where min 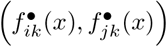 captures the minimum of the two functions evaluated at *x* and *μ*(*x*) is the counting measure. We refer to the kernel function as the cAUC kernel as it computes the minimal common area under the two individuals’ duplication and deletion profile curves. The cAUC kernel function is a valid kernel as shown in S1 Appendix.

The intuition of the cAUC kernel is to quantify similarity using the length of overlapping CNVs between two individuals, with dosage information of the two overlapping CNVs determining how the overlapping length is scaled. The similarity between CNVs of different types (i.e., duplication vs. deletion) is 0, and the similarity between CNVs of the same type depends on the copy number values and the dosage transform function *a*^•^(*DS*). For legal choices of *a*^•^(*DS*), the similarity between a shared duplication (or deletion) event of larger magnitude will be higher than the similarity between a duplication of smaller magnitude, while the minimum operator enforces that the overlapping length is scaled by the CNV of smaller magnitude in a pair with different magnitudes.

Legal choices of *a*^•^(*DS*) will upweight the contribution from similar CNVs of greater magnitude in duplication or deletion, which are often more rare and have higher impact. As proposed in the Discussion section, the family of dosage transform functions *a*^•^(*DS*) = |*DS* − 2|^*d*^ provides a spectrum of weighting schemes, with *d* < 1 down-weighting and *d* > 1 upweighting the contribution of higher magnitude CNVs. Across copy number data of varying types and varying sample-level characteristics, the *a*^•^(·) dosage transform function allows for flexible scaling of dosage to appropriately customize the cAUC measure of similarity.

#### Association test

The association between phenotype and CNVs is examined by testing the hypothesis *H*_0_ : *h*(·) = 0. To do so, we define the vector of subject-specific CNV effects *H* = (*h*(*Z*_1_), ⋯, *h*(*Z_n_*)) and treat *H* as random effects which follow *N*(0, *τ***K**), where *τ* ≥ 0 is a variance component and **K** is a *n* × *n* kernel matrix with its (*i, j*)th entry being *K*(*Z_i_, Z_j_*). Following Liu et al. [34] [35], testing *H*_0_ : *h*(·) = 0 is equivalent to testing *τ* = 0 under a generalized linear mixed model. As in [7] [6], we use a score-based test, which has the form of

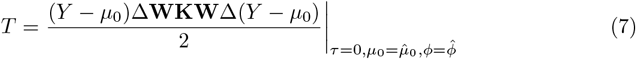

where *Y* is *n* × 1 vector of responses; *μ*_0_ = *E*(*Y*) under *H*_0_; *ϕ* is a dispersion factor parameterizing the variance of *Y*; Δ ∈ ℝ^*n*×*n*^ is a diagonal matrix with its *i*th diagonal element being *δ_i_* = 1/*g*′(*μ_i_*); **W** ∈ ℝ^*n*×*n*^ is a diagonal weight matrix with its *i*th diagonal element being 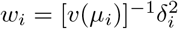 where *v*(·) comes from Var(*Y_i_*) = *v*(*μ_i_*)*ϕ* per the exponential dispersion family of probability density functions. The score statistic asymptotically follows a weighted chi-square distribution [34] [35]. Recently, Chen et al. [12] derived the corresponding small-sample distribution, which is used to calculate the p-value in this work.

## Supporting information

**S1 Table. Gene-level CONCUR tests on genes in pathway hsa01040.**

**S1 Appendix. Proof of symmetry and positive semi-definiteness of cAUC kernel.**

